# Oxidative Stress Mediates UVC-Induced Increases in Epidermal Autofluorescence of C57 Mouse Ears

**DOI:** 10.1101/298000

**Authors:** Mingchao Zhang, Dhruba Tara Maharjan, Yujia Li, Weihai Ying

## Abstract

Our recent study has reported that UV-induced epidermal autofluorescence (AF) can be used as a novel biomarker for predicting UV-induced skin damage, which is originated from UV-induced, cysteine protease-mediated keratin 1 degradation. A key question regarding these findings is: Does oxidative stress play a significant role in the UV-induced epidermal AF and keratin 1 proteolysis? In our current study, we administered the widely used antioxidant N-acetyl cysteine (NAC) into the skin of mouse ears to test our hypothesis that oxidative stress mediates UV-induced increases in the epidermal AF and keratin 1 degradation. Our study has shown that NAC administration can significantly attenuate the UVC-induced AF increases. The NAC administration can also significantly decrease the UVC-induced keratin 1 degradation. Collectively, our findings have indicated that the oxidative stress induced by UVC is causative to the UVC-induced increases in epidermal AF and keratin 1 proteolysis. Moreover, since oxidative stress is significantly increased in multiple regions of the body in several major diseases, the oxidative stress-induced increases in epidermal AF may become a novel biomarker for diagnosis of major diseases.

## Introduction

Human autofluorescence (AF) of skin or blood has been used for non-invasive diagnosis of diabetes (13) and cancer (19). Our recent study has reported that UV-induced epidermal AF can be used as a novel biomarker for predicting UV-induced skin damage (11). Our study has also suggested that the AF increases are originated from UV-induced, cysteine protease-mediated keratin 1 degradation (11): While keratin 1 is an endogenous fluorophore (7), UV-induced, cysteine protease-mediated keratin 1 degradation can lead to increased AF of keratin 1.

Keratins play multiple significant roles in epithelium, including intermediate filament formation (15), cellular signaling (2,10), and inflammatory responses (8,14). Keratin 1 and its heterodimer partner keratin 10 are the major keratins in the suprabasal keratinocytes of epidemis (6,12,18), which is a hallmarker for keratinocyte differentiation (20). Cumulating evidence has also suggested new biological functions of keratins, e.g., keratin 1 has been used as a diagnostic tumor marker (10).

Oxidative stress plays critical roles in UV-induced skin damage (4,9). A key question regarding our finding of UV-induced keratin 1 proteolysis is: Does oxidative stress play significant role in the UV-induced keratin 1 proteolysis? In our current study, we applied the widely used antioxidant NAC to test our hypothesis that oxidative stress mediates UV-induced increases in the epidermal AF and keratin 1 degradation. Our study has shown that NAC administration can significantly attenuate both UVC-induced AF increases and UVC-induced keratin 1 degradation, which has provided evidence supporting our hypothesis.

## Materials and Methods

All chemicals were purchased from Sigma (St. Louis, MO, USA) except where noted.

### Animal Studies

Male C57BL/6Slac mice, ICR mice, and BALB/cASlac-nu nude mice of SPF grade were purchased from SLRC Laboratory (Shanghai, China). All of the animal protocols were approved by the Animal Study Committee of the School of Biomedical Engineering, Shanghai Jiao Tong University.

### Exposures of UV radiation

As described previously (11), an UVC lamp (TUV 25W/G25 T8, Philips, Hamburg, Germany) was used as the UV sources in our experiments. C57BL/6Slac mice at the weight between 18-35 g were used for UVC treatment. After the mice were briefly anesthetized with 3.5% (w/v) chloral hydrate (1 ml/100g), the ears of the mice were exposed to UV lamps. The power density of UVC was 0.55±0.05 mW/cm^2^, measured by a UVC detector (ST-512, UVC, SENTRY OPTRONICS CORP., Taiwan, China).

### Imaging of epidermal AF

As described previously (11), the epidermal AF of the ears of the mice was imaged by a two-photon fluorescence microscope (A1 plus, Nikon Instech Co., Ltd., Tokyo, Japan), with the excitation wavelength of 488 nm and the emission wavelength of 500 - 530 nm. The AF was quantified by the following approach: Sixteen spots with the size of approximately 100 × 100 µm^2^ on the scanned images were selected randomly. After the average AF intensities of each layer were calculated, the sum of the average AF intensities of all layers of each spot was calculated, which is defined as the AF intensity of each spot. If the value of average AF intensity of certain layer is below 45, the AF signal of the layer is deemed background noise, which is not counted into the sum.

### Western blot assays

As described previously (11), the lysates of the skin were centrifuged at 12,000 *g* for 20 min at 4°C. The protein concentrations of the samples were quantified using BCA Protein Assay Kit (Pierce Biotechonology, Rockford, IL, USA). As described previously(17), a total of 50 μg of total protein was electrophoresed through a 10% SDS-polyacrylamide gel, which were then electrotransferred to 0.45 μm nitrocellulose membranes (Millipore, CA, USA). The blots were incubated with a monoclonal Anti-Cytokeratin 1 (ab185628, Abcam, Cambridge, UK) (1:4000 dilution) or actin (1:1000, sc-58673, Santa Cruz Biotechnology, Inc., Dallas, TX, USA) with 0.05% BSA overnight at 4°C, then incubated with HRP conjugated Goat Anti-Rabbit IgG (H+L) (1:4000, Jackson ImmunoResearch, PA, USA) or HRP conjugated Goat Anti-mouse IgG (1:2000, HA1006, HuaBio, Zhejiang Province, China). An ECL detection system (Thermo Scientific, Pierce, IL, USA) was used to detect the protein signals. The intensities of the bands were quantified by densitometry using Image J.

### Laser-based delivery of NAC into mouse skin

Male C57BL/6SlacMice were briefly anesthetized with 3.5% (w/v) chloral hydrate (1 ml / 100 g). After exposure to laser, the ears of the mouse were administered with NAC. Stock solutions of 100 mM and 250 mM NAC were prepared by a mixture of PBS and equal volume of glycerol.

### Statistical analyses

All data are presented as mean +SEM. Data were assessed by one-way ANOVA, followed by Student – Newman - Keuls *post hoc* test. *P* values less than 0.05 were considered statistically significant.

## Results

We found that exposures of the skin of mouse ears to 0.66 J/cm^2^ UVC led to increases in the skin AF of the ears of C57 mice, assessed at 0, 1, 6 or 24 hrs after the UVC exposures (Fig. 1A). Quantifications of the AF showed that each dose of UVC induced significant increases in the epidermal AF of the ears of C57 mice (Fig. 1B).

**Figure 1.**
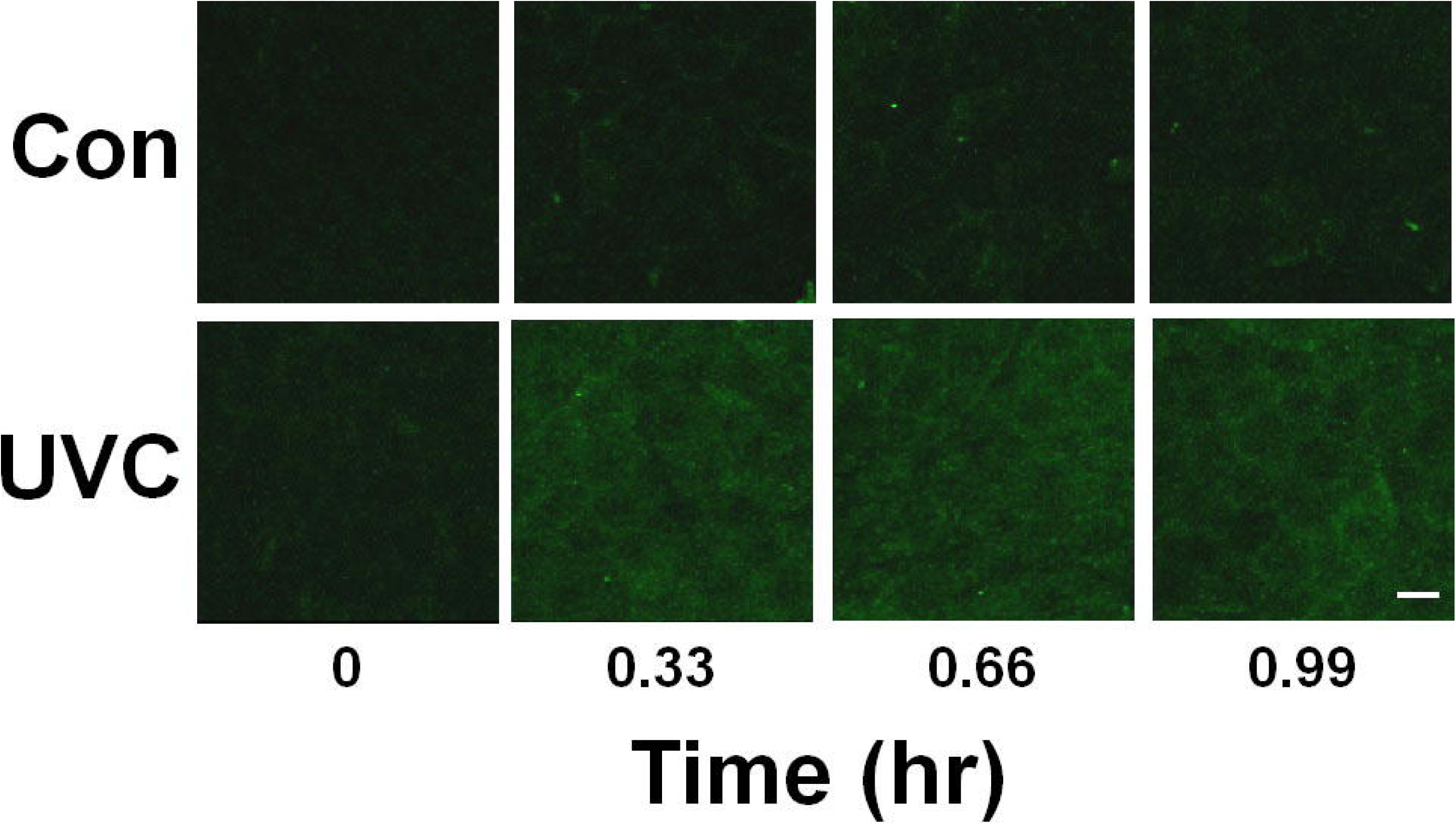
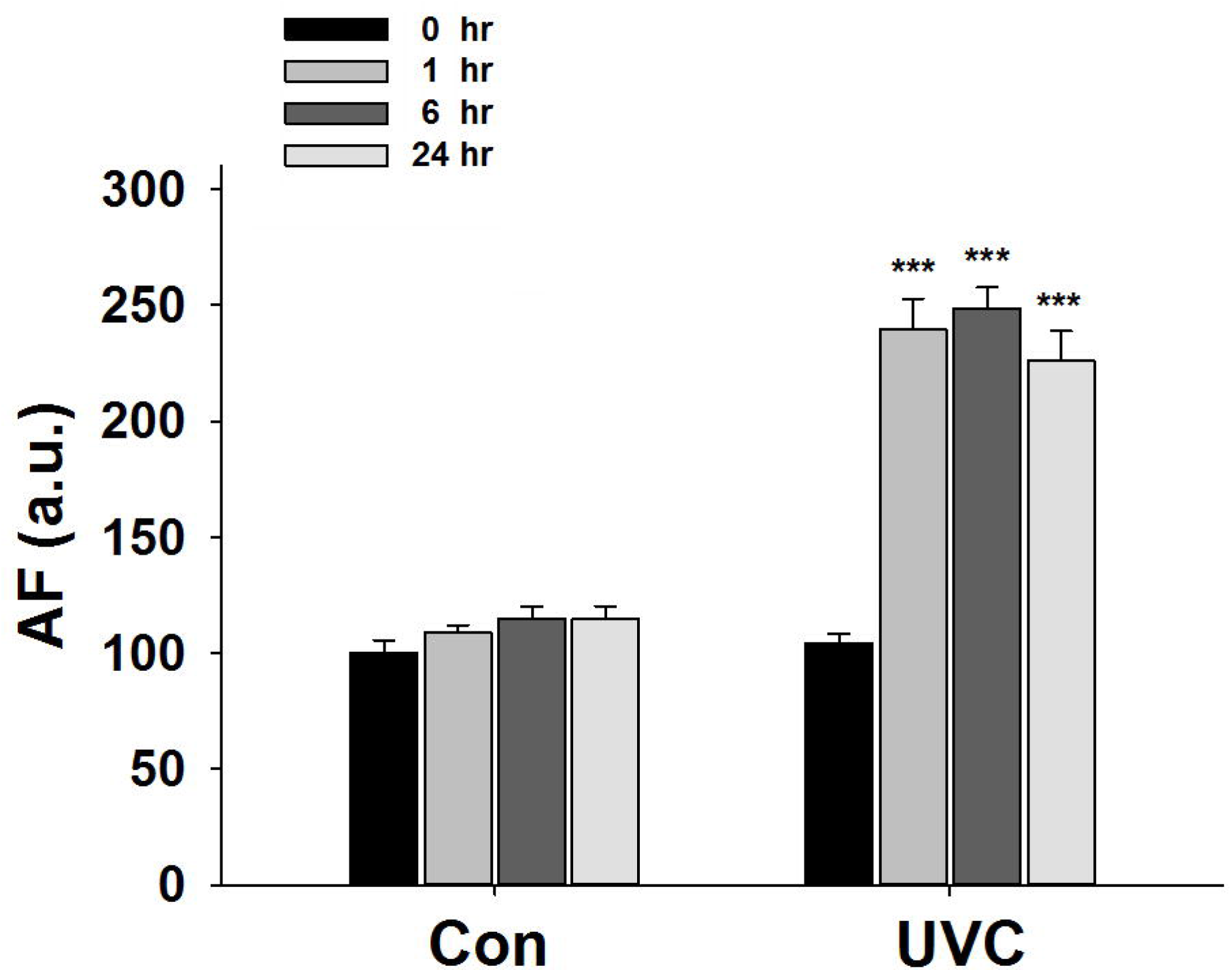
UVC induced increases in the epidermal autofluorescence of C57 mouse ears. (A) Exposures of the skin to 0.66 J/cm^2^ UVC led to increases in the skin AF of the ears of C57 mice, assessed at 0, 1, 6 or 24 hrs after the UVC exposures. Excitation wavelength = 488 nm and emission wavelength = 500 - 530 nm. Scale bar = 20 µm. (B) Quantifications of the AF showed that each dose of UVC induced significant increases in the skin AF of the ears of C57 mice. N = 10 – 13. *, *P* < 0.05; ***, *P* < 0.001.

Our AF imaging study showed that administration of 5 or 12.5 mg/cm^2^ NAC led to decreases in the 0.66 J/cm^2^ UVC-induced epidermal AF, assessed at 1 hrs after the UVC exposures (Fig. 2A). Quantifications of the AF images showed that NAC dose-dependently attenuated the UVC-induced epidermal AF (Fig. 2B).

**Figure 2.**
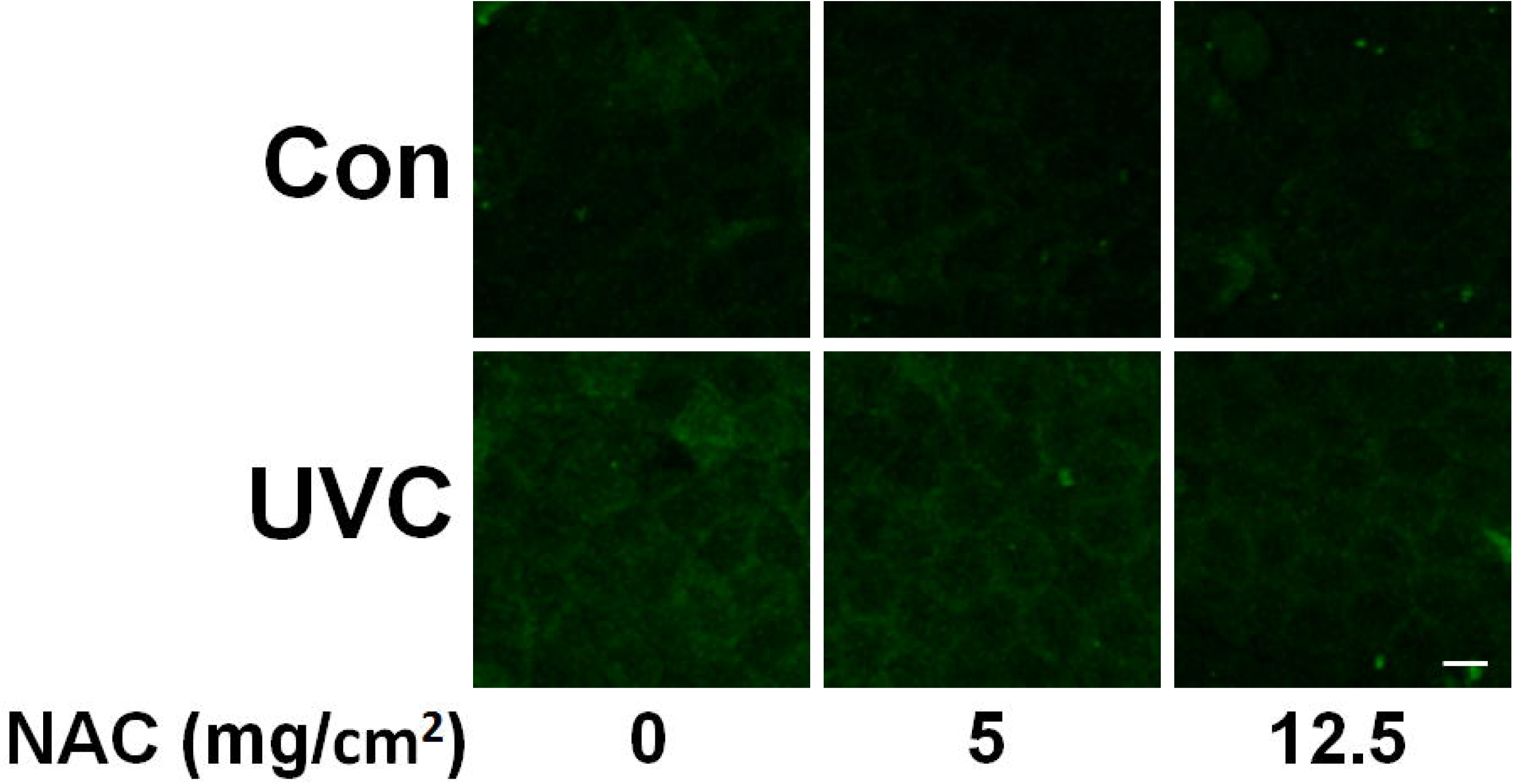
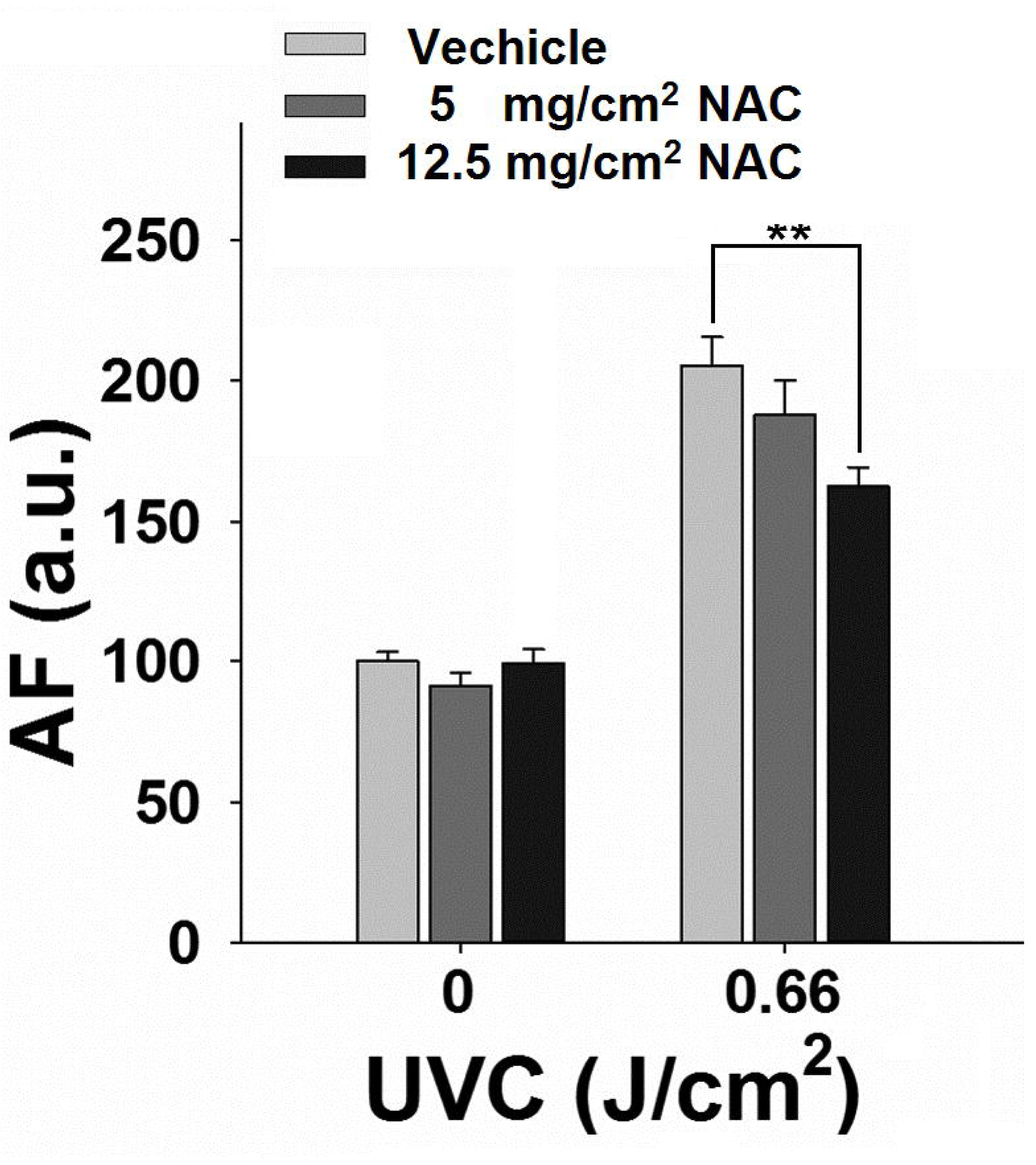
NAC administration decreased UVC-induced epidermal autofluorescence of C57 mouse ears. (A) AF imaging study showed that administration of NAC led to decreases in the 0.66 J/cm^2^ UVC-induced epidermal AF, assessed at 1 hrs after the UVC exposures. (B) Quantifications of the AF images showed that NAC significantly attenuated the UVC-induced epidermal AF. N = 12 - 15. **, *P* < 0.01.

Our Western blot assays showed that administration of attenuated the UVC (0.66 J/cm^2^)-induced degradation of keratin 1, assessed at 1 hrs after the UVC exposures (Fig. 3A and 3B). In contrast, administration with NAC led to prevention of the UVC-induced degradation of keratin 1 (Fig. 3A and 3B).

**Figure 3.**
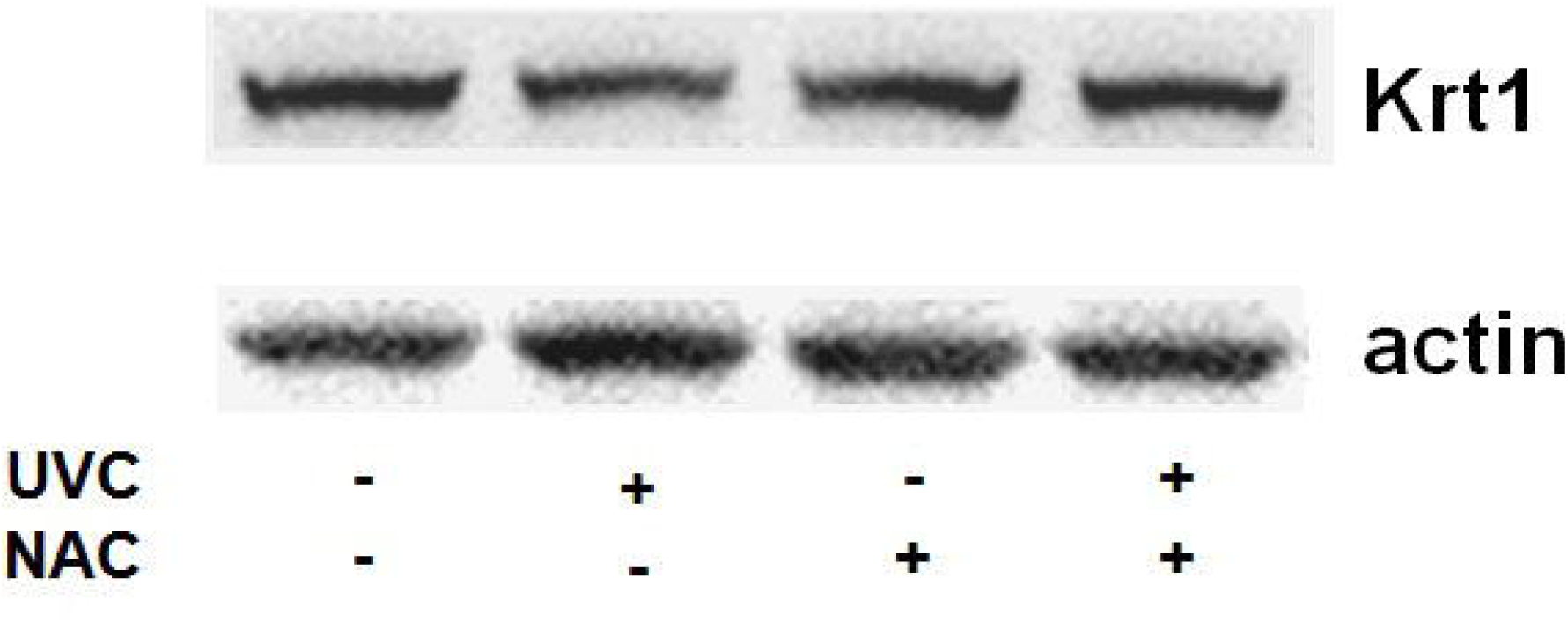
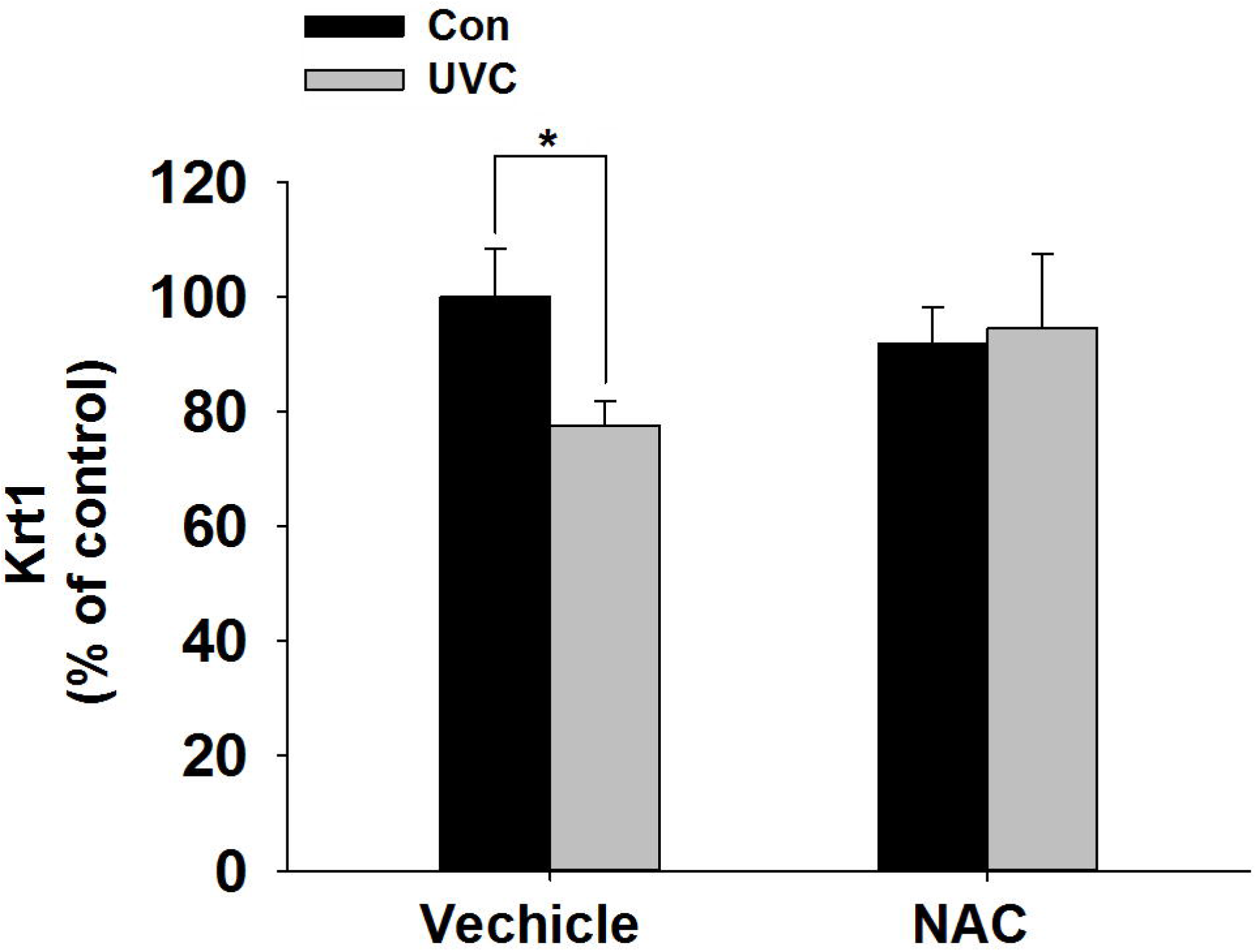
NAC administration decreased UVC-induced keratin 1 degradation of C57 mouse ears. (A) Western blot assays showed that administration of NAC led to decreases in the 0.66 J/cm^2^ UVC-induced degradation of keratin 1, assessed at 1 hrs after the UVC exposures. (B) Quantifications of the Western blot showed that NAC significantly attenuated the UVC-induced degradation of keratin 1,**, *P* < 0.01. N = 12 - 15.

## Discussion

The major findings of our study include: First, NAC administration can significantly attenuate UVC-induced increases in the epidermal AF of C57 mouse ears; and second, NAC administration can significantly reduce the UVC-induced keratin 1 degradation of C57 mouse ears. Collectively, these results support our hypothesis that oxidative stress mediates UVC-induced increases in the epidermal AF and keratin 1 degradation, which have indicated that the oxidative stress induced by UVC is causative to the UVC-induced increases in the epidermal AF and keratin 1 proteolysis.

Our recent study has reported that UV-induced epidermal AF can be used as a novel biomarker for predicting UV-induced skin damage (11). Our study has also suggested that the AF increases are originated from UV-induced, cysteine protease-mediated keratin 1 degradation. Because oxidative stress plays critical roles in UV-induced skin damage (4,9), it is necessary to elucidate the roles of the UV-induced AF increases and keratin 1 proteolysis. Our current study has suggested that oxidative stress mediates the UV-induced AF increases and keratin 1 proteolysis. Since our previous study has suggested that keratin 1 proteolysis is an important mechanism underlying UV-induced skin damage (11), our finding regarding the role of oxidative stress in UV-induced keratin 1 degradation has suggested an mechanism underlying the role of oxidative stress in UV-induced skin damage: Oxidative stress could mediate UV-induced skin damage partially by producing keratin 1 degradation. It is warranted to determine the mechanisms underlying the roles of ROS in the UV-induced AF increases and keratin 1 proteolysis. We propose that the UV-induced ROS may produce these effects by activating the enzyme that produces the keratin 1 proteolysis, or by altering the structure of keratin 1 which makes the keratin 1 more susceptible to the proteolytic effects of the enzyme.

A number of studies have shown that oxidative stress is significantly increased in multiple regions of the body in multiple major diseases, such as cerebral ischemia (3,5) and myocardial infarction (1,16). It is possible that the oxidative stress may induce increases in epidermal AF of the patients of these diseases, which may become a novel marker for non-invasive and efficient diagnosis of major diseases. Future studies are warranted to determine if there are increases in oxidative stress-induced epidermal AF in the body of certain major diseases.

## Acknowledgments

The authors would like to acknowledge the financial support by a Major Research Grant from the Scientific Committee of Shanghai Municipality #16JC1400502 (to W.Y.) and Chinese National Natural Science Foundation Grant #81271305 (to W. Y.).

